# Origami: Single-cell oriented 3D shape dynamics of folding epithelia from fluorescence microscopy images

**DOI:** 10.1101/2021.05.13.443974

**Authors:** Tania Mendonca, Ana A. Jones, Jose M. Pozo, Sarah Baxendale, Tanya T. Whitfield, Alejandro F. Frangi

## Abstract

A common feature of morphogenesis is the formation of three-dimensional structures from the folding of two-dimensional epithelial sheets aided by spatio-temporal cell shape changes at the cellular-level. Studying cell shape dynamics and polarised processes that underpin them, requires orienting cells within the epithelial sheet. In epithelia with highly curved surfaces, assigning cell orientation can be difficult to automate *in silico*. We present ‘Origami’, a MATLAB-based image analysis pipeline to compute oriented cell shape-features. Our automated method accurately computed cell orientation in regions with opposing curvature in synthetic epithelia and fluorescence images of zebrafish embryos. As proof of concept, we identified different cell shape signatures in the developing zebrafish inner ear, where the epithelium deforms in opposite orientations to form different structures. Origami is designed to be user-friendly and is generally applicable to fluorescence images of curved epithelia.

## Introduction

Complex morphologies across taxa and tissue types are generated through the deformation of epithelial sheets [1–3]. In the embryo, many developing epithelia form highly curved surfaces. Epithelial folding processes are driven by polarised mechanical forces and involve three-dimensional changes in shape at the cellular level [4,5]. Fluorescence imaging techniques have made it possible to follow such shape changes at cellular resolution, *in vivo* and in real-time [6–8]. These imaging advances have consequently driven the development of tools to quantify epithelial dynamics, especially cell shape changes.

Many image analysis tools measuring cell shape change have been limited to two-dimensional [9–12] or quasi-3D fluorescence microscopy data [13]. Extending these measurements to 3D has been aided by the development of membrane-based 3D segmentation methods such as ACME [14], RACE [15], 3DMMS [16], CellProfiler 3.0 [17], and more recently, deep-learning-based methods [18–21]. Some image analysis tools, such as CellProfiler 3.0 [17], MorphoGraphX [22] and ShapeMetrics [23], provide pipelines to compute unoriented cell shape features. However, orienting the 3D-segmented cells along biologically relevant axes to quantify directionally variant shape features is still a challenging problem that has so far not seen a generalised solution.

Orienting cells relative to the known overall polarity of the epithelial sheet is critical, as cell polarised biomechanical processes drive cell shape changes; constriction or expansion can occur along either the apical [24,25] or baso-lateral [26] cell surfaces and can be detected by any skew in mass distribution within a cell along an apico-basal axis of symmetry. Epithelial folding may be initiated or influenced by cell proliferation, cell death, cytoskeletal remodelling, or changes in cell surface properties [27,28]. These mechanisms can lead to changes in cell shape features, including cell height and width, volume, surface area and sphericity.

Cell orientation or polarity can be defined along the plane of the epithelium (planar cell polarity) or perpendicular to the epithelial plane, along the apico-basal axis of the cell. Existing automated methods for assigning polarity often rely on additional biochemical markers for polarity [29–31]. Including such additional markers in fluorescence imaging experiments increases the time taken to generate each image, potentially leading to phototoxicity, and the resulting larger volume of image data makes analysis computationally expensive. Moreover, producing the required animals carrying multiple transgenes for live imaging can be challenging and costly. Some image analysis methods orient cells by fitting polynomial functions, often ellipsoids, to estimate the surface of the specimen — for example, entire embryos [15] or blastoderms [32] undergoing morphogenesis — and draw normal vectors to the fitted folding surface at each cell. These methods are specific to the geometry of the specimen and are unsuitable for analysing complex folded topologies at advanced morphologic developmental stages. A third method uses known features of cell shape to assign orientation, for example by applying principal component analysis (PCA) to compute the apico-basal direction in columnar cells in EDGE4D [33] and the anterior-posterior axis in zebrafish lateral line primordia using landmark-based geometric morphometrics [31], or orienting cells along their long axis in the zebrafish optic cup as in LongAxis [34]. These strategies will be applicable only if a shape feature is known.

We introduce a new automated tool, Origami, for extracting shape features oriented along the apico-basal axis by reconstructing the epithelial surface using a triangular mesh (Fig 1). Origami applies to a wide range of geometries of specimens undergoing morphogenesis and computes the apico-basal axis of the epithelial sheet for known epithelial organisation without requiring additional labels for polarity . We showcase the versatility of our method using data from an assortment of structures at a range of developmental stages within the otic vesicle (developing inner ear) of zebrafish embryos.

**Fig 1:**
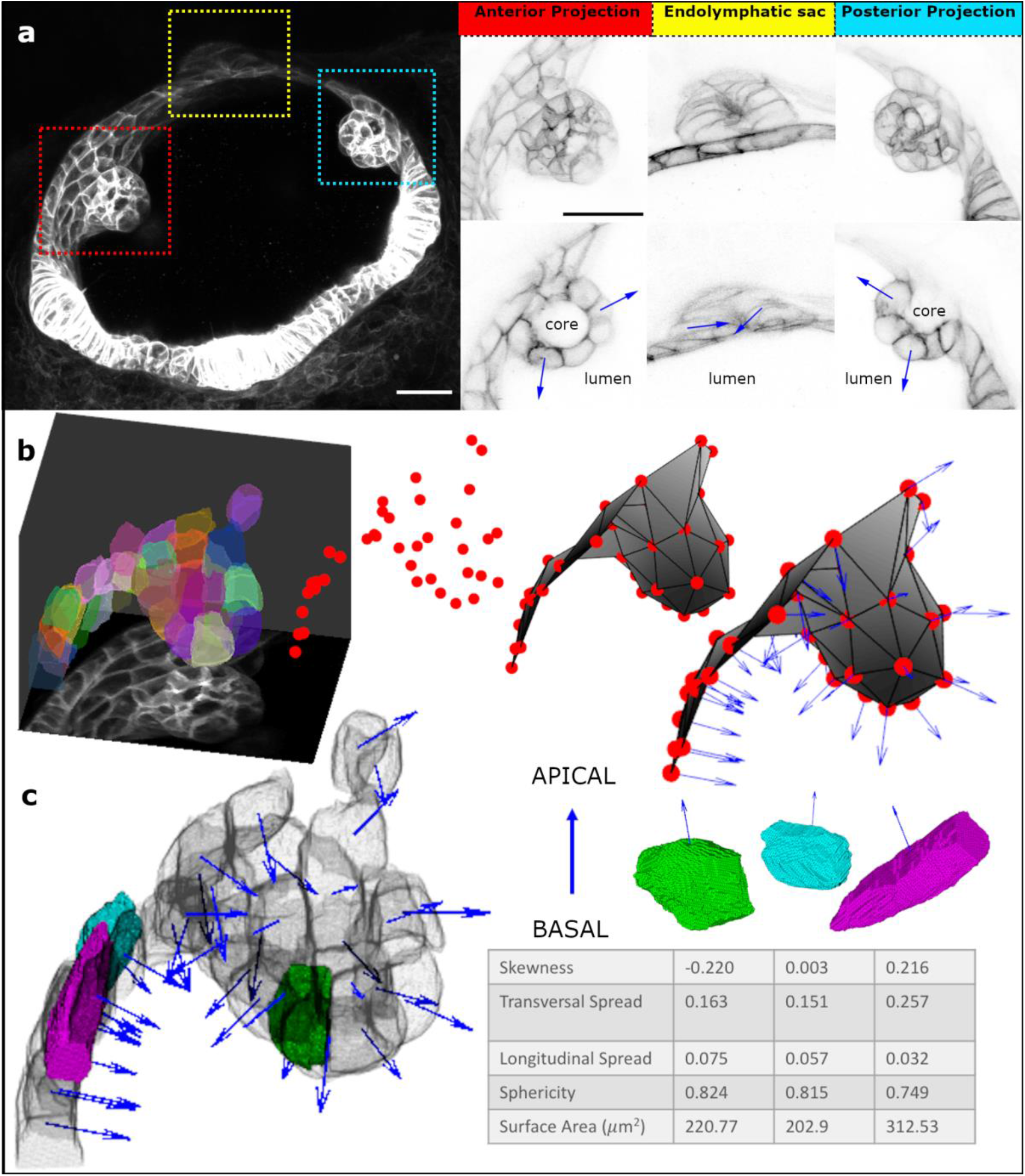
Origami Image Analysis Pipeline. **a**. Airyscan confocal fluorescence micrograph (maximum intensity projection (MIP) of 35 z-slices) of the developing zebrafish otic vesicle at 51.5 hours post fertilisation. Red box – anterior projection; yellow box – endolymphatic sac; cyan box – posterior projection. The ROIs are expanded alongside – top row MIPs, and bottom row single slices. Scale bars: 20 µm. Blue arrows mark the direction of apicobasal polarity (pointing towards the apical side). **b**. Polarity assignment on segmented data; ROI surrounding the anterior projection was segmented (here overlaid on the MIP) using ACME, centroids were generated for each segmented cell and a triangular surface mesh was produced from these centroids. Normal vectors (blue arrows) to this surface mesh represent the apico-basal axis. **c**. Cell shape features were computed concerning the assigned apico-basal axis; here, three example cells are highlighted, alongside a 3D rendering showing their position in the anterior projection and the corresponding shape metrics in a table.

## Design and Implementation

The Origami pipeline is preceded by a membrane-based segmentation step. For this, we employed the open-source ACME segmentation software [14]. The segmented data are subjected to two main operations within Origami; epithelial orientation assignment (Fig 1b) and extraction of shape features (Fig 1c).

### Assigning orientation to individual cells

To compute directionally variant cell shape features, such as skewness (asymmetry in cell mass), and longitudinal and transversal spread, segmented cells need to be oriented in 3D space along a biologically relevant axis – we chose the known apico-basal axis of the cell. The folding epithelium was reconstructed *in silico* as a thin ‘crust’ – an open surface mesh that triangulates the centroids of the segmented cells in 3D space using the Crust algorithm [35,36] (Fig 1b). The Crust method computes a surface mesh from unorganised points – cell centroids in our case, using the Voronoi diagram of the cell centroids.

Following this, our automated method corrects imperfections in the estimated surface mesh that can cause polarity assignment errors. The mesh is refined by removing duplications (in vertices or triangular faces computed) and any self-intersecting triangular faces. Non-manifold edges, that is, those edges shared by more than two triangular faces, are re-meshed as a manifold mesh using the ball-pivoting algorithm [37,38].

The triangular faces of the refined mesh are ordered in the same direction, and so by applying the right-hand rule when generating normal vectors to the surface mesh, these vectors all point to the same side of the surface (Fig 1b). In the developing zebrafish otic vesicle, the otic epithelium shows an apico-basal polarity, with the apical surface facing the fluid-filled lumen of the vesicle [2,8,39]. We used this prior knowledge to assign apico-basal polarity as a vector pointing to the side of the curved surface mesh that corresponds to the apical (lumenal) side of the epithelium.

Under-segmentation can cause missing regions or unwanted holes in the triangular mesh, introducing errors when ordering the triangular faces. Our pipeline attempts to repair these holes by detecting and then remeshing them where possible. Holes, when detected, are flagged as a warning to users about potential errors in the output. Normal vectors to the reconstructed surface represent the epithelium’s apico-basal axis and are generated for each segmented cell at their centroid position (Fig 1b and c).

### Computing shape features using 3D geometric moments

The shape of an object can be characterised using central geometric moments [40]. Geometric moments are widely used in object recognition and classification problems [41,42] since they (i) are simple to compute, (ii) organise features in orders of increasing detail, and (iii) can be extended to *n* dimensions. Each moment, 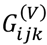,is defined by the integral over the object (in our case, each segmented cell), of the Cartesian coordinates monomial *x*^*i*^*y*^*j*^*z*^*k*^, where *i, j, k* ≥ 0, with the origin of coordinates at the centroid.

In our analysis pipeline, 3D geometric moments were computed from triangular surface meshes generated for each individual segmented cell [43]. In this method, the integral defining the geometric moments of each segmented cell is split into a sum:

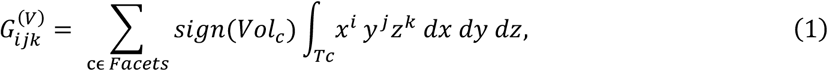

where each tetrahedron T_c_ is defined by a triangle in the surface mesh and the origin (cell centroid). The determinant gives the oriented volume of this tetrahedron,

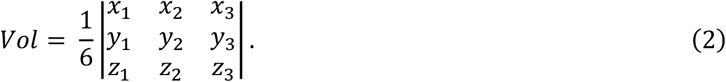

Considering its sign, the determinant allows the algorithm to be applied to shapes of any complexity and topology. Thus, the geometric moments of each segmented cell can be directly calculated from the Cartesian coordinates of the triangular surface mesh vertices.

The geometric moments of low orders have simple, intuitive interpretations. The zero^th^ order moment 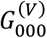 provides the volume of the object, here an individual cell. For central moments, the first order moments are trivially null: 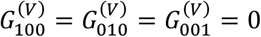. The second-order moments correspond to the spread (covariance tensor) of the distribution. So, the projection of the mass of each cell along the corresponding polarity vector represents the ‘spread’ as variance in mass ‘longitudinally’ (along the apico-basal axis) and ‘transversally’ (along the epithelial plane). This allowed us to identify if cells were more or less columnar or squamous in shape. The third-order moments represent ‘skewness’, which is the deviation from symmetry. In our pipeline, skewness was measured along the polarity vector in the apico-basal direction, with positive skewness values indicating apical cell constriction and/or basal relaxation and negative values indicating basal cell constriction and/or apical expansion. A value of zero indicated no skew. Additionally, the sphericity of each cell was computed as the ratio of the cell surface area to the surface area of a sphere with the same volume as the cell [44], from 0 for a highly irregularly-shaped cell to 1 for a perfect sphere.

## Results

### Evaluation of Computed Cell Orientation

To evaluate computed cell orientation, we generated 3D synthetic data representing curved, folding epithelia with varying degrees of curvature and height of folded peak in two opposing directions (Supplementary Materials and Fig 2a). To reflect real-world *in vivo* fluorescence imaging conditions, these synthetic data were corrupted with three incremental levels of Gaussian and Poisson noise (Supplementary Materials and Fig 2a). Using the synthetic data, two types of error in cell orientation were assessed: (1) an orientation flipping error, measured as the percentage of polarity vectors pointing in the opposite direction to the polarity ground truth (Supplementary Materials), and (2) direction accuracy, measured as the mean deviation angle between the polarity vectors produced by Origami, correctly oriented, and the polarity ground truth.

**Fig 2:**
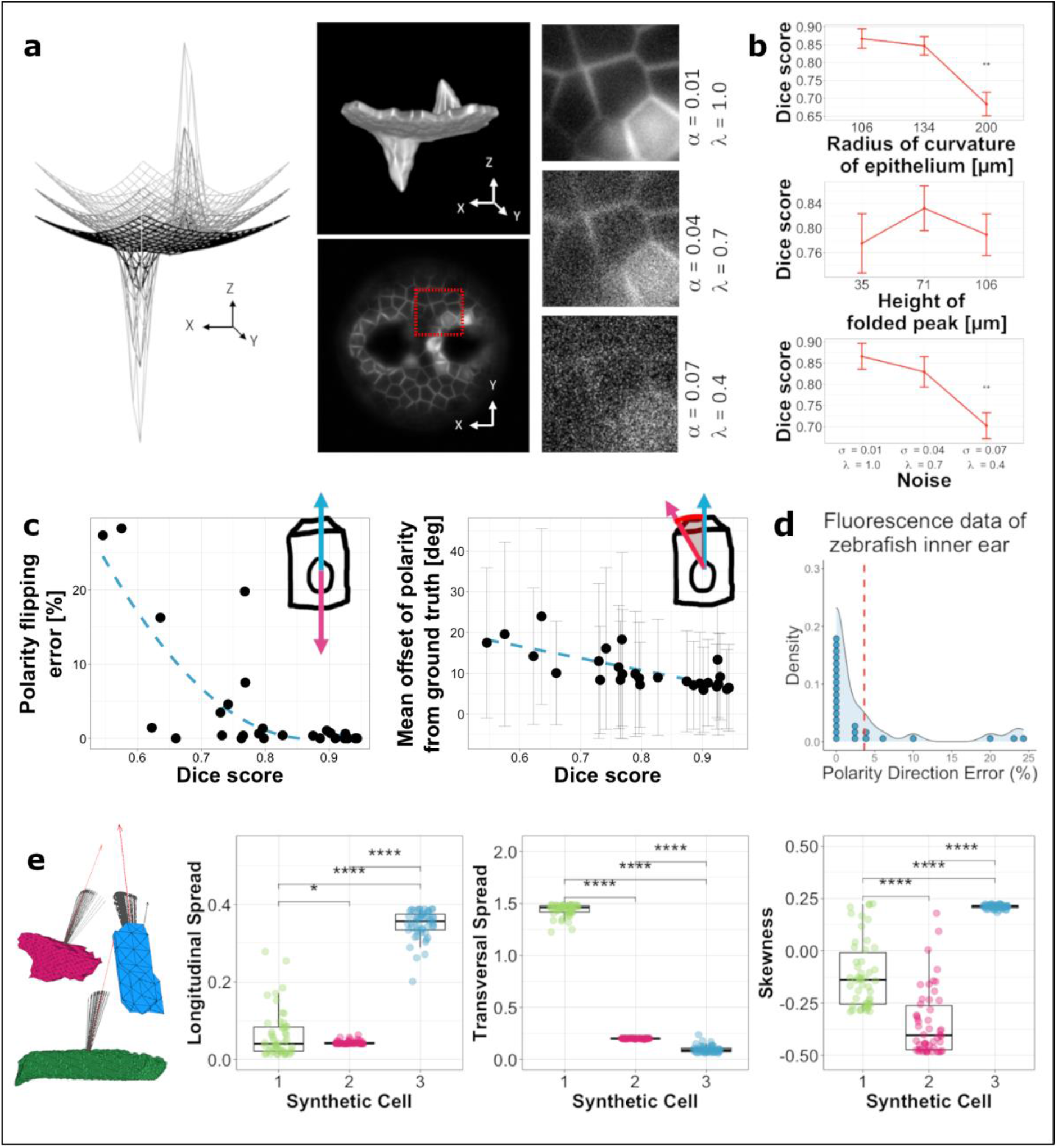
Assessment of polarity assignment. **a**. Surface meshes of synthetic epithelia for validating the Origami analysis pipeline. Alongside, 3D rendering of one of the synthetic epithelia (top) and a single 2D slice through it (bottom). Each image volume was corrupted with three levels of noise. **b**. The relationship between surface geometry/ noise and segmentation quality. Error bars represent the standard deviation. Tukey’s pairwise comparisons with significant values depicted with asterisks; Dice score at radius of curvature of 200 µm (1000 pixels) compared to that at 106 µm (530 pixels) – p = 0.0004, Dice score at largest noise level compared to the lowest – p = 0.0045. **c**. Effect of segmentation quality on errors in orientation flipping (left) and direction offset in the computed polarity vectors (error bars in grey represent standard deviation). Dashed lines represent quadratic and linear fit to data respectively. **d**. Probability density of errors in polarity direction in real fluorescence data from zebrafish embryos. Each dot represents the percentage error from a 3D segmented volume (n = 27; total of 949 segmented cells across all the images). The dashed line shows the mean error in the dataset (<4%). **e**. Sensitivity of cell shape metrics to errors in polarity orientation. Data points in the graphs are depicted with the same colour as the corresponding synthetic cell alongside. Tukey’s pairwise comparisons with significant values depicted with asterisks; Longitudinal spread: 1-2 p = 0.039, 1-3 p < 0.0001, 2-3 p < 0.0001; Transversal Spread: 1-2 p < 0.0001, 1-3 p < 0.0001, 2-3 p < 0.0001; Skewness: 1-2 p < 0.0001, 1-3 p < 0.0001, 2-3 p < 0.0001.

Of the two aspects of surface geometry analysed, height of folded peak (in two opposing directions), did not contribute significantly to orientation flipping errors (Linear Regression; *p* = 0.86, *R*^2^ = -0.04). However, a larger radius of curvature of epithelium (a flatter epithelial sheet), did correlate with orientation flipping errors – albeit with a small effect of 0.08% increase for every 1 µm (5 pixels) increase in radius of curvature (Linear Regression; p = 0.042, R^2^ = 0.12, effect), and a lower quality of segmentation output from ACME (Linear Regression *p* < 0.001, *R*^2^= 0.46; Fig 2b) computed as a Dice score. This meant a 0.2% reduction in Dice score for every 1 µm (5 pixels) increase in the radius of curvature. This correlation may be attributed to the reduced ability of ACME to segment flat, squamous cells in an epithelium oriented mostly along the lateral (XY) plane in data with anisotropic voxel resolution (here modelled using an anisotropic point spread function (PSF)). We found a correlation between noise applied to the synthetic images and errors in both polarity orientation flipping (ANOVA: *p* ≈ 0.001; Tukey’s contrasts showed 11.3% increase in errors at highest noise level compared with the lowest noise level applied – *p* = 0.0039) and segmentation output (ANOVA: *p* < 0.01; Tukey’s contrasts showed 16.3% reduction in Dice score at highest noise level from the lowest noise level applied – *p* = 0.0045). Segmentation quality, in turn, influenced polarity orientation flipping, with errors below 1.5% at Dice scores above 0.8, but increasing with further decrease in Dice scores (Polynomial Regression; first-order: *p* < 0.001, Effect size = -28.78; second-order: *p* < 0.01, Effect size = 16.26; Fig 2c). Comparisons of many available segmentation algorithms when validating with fluorescent images from non-folded structures such as early-stage nematode embryos [16] or plant roots [18] have been shown to give Dice scores above 80%, suggesting a good performance under real experimental conditions.

Quantitative direction accuracy was evaluated in the synthetic data, for which, in contrast to data from real fluorescence images, a reliable ground truth could be generated from the known underlying surface functions. Compared to the polarity ground truth data, an overall offset of 10.6° ± 15.5° (mean ± std) was measured from our entire synthetic dataset. Just as for the polarity orientation flipping error, height of folded peak did not influence polarity direction accuracy (Linear Regression; *p* = 0.39, *R*^2^ = -0.01), but there was a small effect of curvature of the epithelium with an additional 0.06° offset for every 1 µm (5 pixels) increase in radius of curvature of the epithelium (Linear Regression; *p* = 0.005, *R*^*2*^ = 0.24). At the highest level of noise applied, errors in polarity orientation had a 6.6° greater offset than at the lowest noise level applied (Tukey’s contrasts; *p* = 0.003). There was also a negative linear effect of segmentation quality with a 2.9° offset predicted for every 10% reduction in Dice score (Linear Regression; *p* < 0.0001, *R*^2^ = 0.53; Fig 2c).

We further tested the effect of such errors in direction accuracy on the oriented shape metrics computed by applying noise in orientation—with a mean equal to the measured mean error above —to polarity vectors of three example cells showing extreme shape features from the synthetic dataset and computed oriented shape metrics for each new displaced polarity vector (*n* = 50; Fig 2e). The resulting computed shape metrics could still successfully differentiate between the three cells, showing that direction accuracy errors (excluding flipping errors) should not adversely affect the shape metrics computed. On the other hand, orientation flipping errors will affect shape metrics, but as shown above, these errors are predicted to be small for a well-segmented image volume and can be easily identified by visual inspection and corrected if needed using the Origami pipeline.

Additionally, orientation flipping errors were quantified from real light-sheet fluorescence microscopy data from structures in the developing zebrafish otic vesicle (Figs. 1, 3). For this, cells assigned orientation in the opposite direction to the apico-basal polarity were identified by visual assessment in the Origami pipeline, showing errors in 3.65% of *n* = 949 cells analysed (Fig 2d).

**Fig 3:**
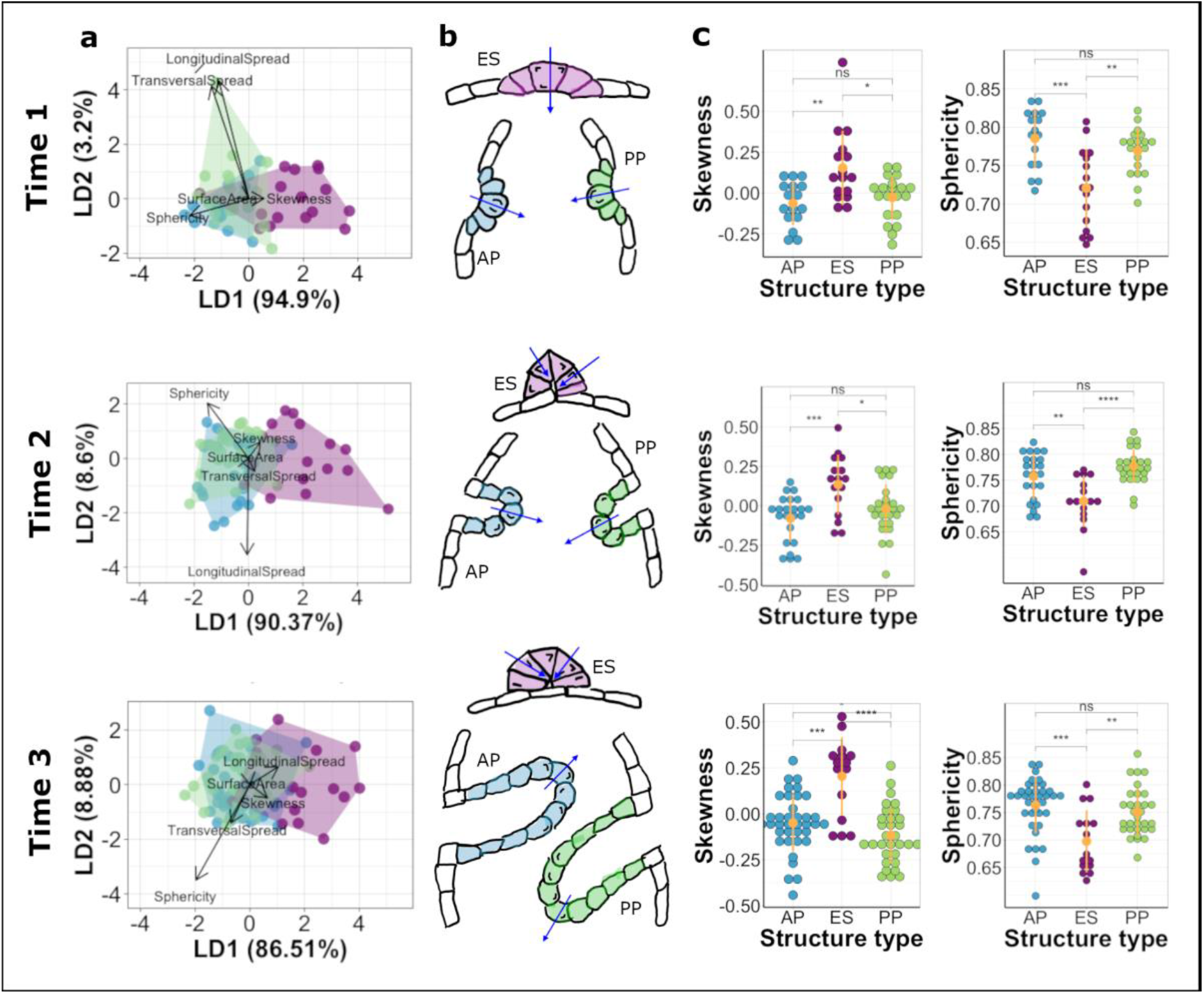
Comparison of shape dynamics in developing structures of the zebrafish inner ear. Rows represent each time point analysed. Data in blue represent cells from AP, green represent cells from PP and magenta represent ES. **a**. Linear discriminate analysis (LDA) biplots illustrate multivariate clustering of data – data from AP and PP show considerable overlap indicating similar shape signatures while data from ES show less overlap with the former. **b**. Schematic illustrations of cell shape signatures at the time points analysed showing cells in the ES having skew in the opposite direction to those in the projections and having less rounded shapes. Arrows indicate apico-basal polarity. **c**. Plots showing differences in skewness and sphericity between the structures at the time points analysed. Yellow dots with error lines represent mean and standard deviation for the data. p values for paired comparisons depicted are from Table 1.

## Proof of Principle: Insights Into Cell Shape Dynamics During Epithelial Morphogenesis Within The Zebrafish Inner Ear

To further validate our method, we used Origami to characterise cell shape dynamics involved in the formation of different structures in the otic vesicle of the zebrafish embryo (Figs. 1 and 3). We analysed light-sheet fluorescence image data from the anterior epithelial projection (AP) for the developing semicircular canal system, together with the endolymphatic sac (ES), at three developmental time points: 42.5 hours post fertilisation (hpf) (time point 1), 44.5 hpf (time point 2) and 50.5 hpf (time point 3), using different fish for each time point. We also analysed the posterior epithelial projection (PP), a similar structure to the AP, but which develops later [39], at developmentally equivalent time points to that of the AP (46.5 hpf, 50.5 hpf and 60.5 hpf). The AP and PP are finger-like projections of the epithelium that move into the lumen of the vesicle, with the apical side of the cell on the outside of the curved projection surface [39]. By contrast, the ES forms as an invagination from dorsal otic epithelium, with the constricted apical surfaces of the cells lining the narrow lumen of the resultant short duct [8,45,46]. As the ES is formed through deformation of the epithelial sheet with opposite polarity to that of the epithelial projections, we expect cells in the ES and the projections to show significant differences in cell shape. Conversely, we do not expect significant differences in cell shape between the AP and PP cells, which form equivalent structures in the developing ear.

**Table 1:** Paired comparisons using Wilcoxon rank sum exact test (p values – adjusted using ‘Holm’ correction)

| Time point | AP - ES |  |  | PP - ES |  |  | AP - PP |  |  |
| --- | --- | --- | --- | --- | --- | --- | --- | --- | --- |
|  | 1 | 2 | 3 | 1 | 2 | 3 | 1 | 2 | 3 |
| **Skewness** | **0.009** | **0.002** | **<0.001** | **0.036** | **0.036** | **<0.0001** | 0.42 | 0.286 | **0.047\*** |
| **Sphericity** | **0.002** | **0.012** | **<0.001** | **0.005** | **<0.0001** | **0.007** | 0.149 | 0.228 | 0.07 |
| **Surface Area** | **0.001** | **<0.0001** | **0.032** | **0.001** | **<0.0001** | **0.013** | 0.887 | 0.85 | 0.912 |
| **Transversal Spread** | 0.057 | 0.062 | 1 | 0.357 | 0.062 | 1 | 0.357 | 0.897 | 1 |
| **Longitudinal Spread** | 1 | 0.88 | 0.054 | 1 | 0.81 | 0.102 | 1 | 0.46 | 0.582 |

\*The differences in skewness between cells in the AP and PP at the 3<sup>rd</sup> time point tended towards significance. This might be attributed to differences in the lengths of projections, with cells at the leading end of the projection showing more extreme skewness values while cells along the lateral sides showing less skewed shape.

For each structure, the following shape attributes were computed at the single-cell level: surface area, sphericity, longitudinal spread, transversal spread and skewness. Since volume and surface area show high collinearity within our data (Pearson correlation coefficient = 0.98, 95% confidence intervals = [0.977, 0.984]), cell volume was excluded from further analysis. Although images included cells in the non-folding epithelium around the developing structures of interest, only cells from the folding epithelium were analysed. A multivariate analysis ({MANOVA.RM} package [47] in R [v 4.0.0]) of the dependence of cell shape attributes on the epithelial structure from which they are derived at different time points showed a significant difference between the three structures at the first two developmental times (Wald-type statistic; *p* = 0.035 (resampled *p* = 0.001) at time point 1 and *p* < 0.001 (resampled *p* < 0.001) at time point 2) but not at the final time point analysed (*p* = 0.706 (resampled *p* = 0.038)) for all shape attributes. Post-hoc Tukey’s contrasts indicated that cells in the endolymphatic sac showed significantly different shape dynamics from those of cells in both projections (ES - AP *p* = 0.006, PP - ES *p* = 0.012 at time point 1; ES - AP *p* = 0.0002, PP - ES *p* = 0.0002 at time point 2 but ES - AP *p* = 0.192, PP - ES *p* = 0.116 at time point 3). There was no significant difference in the cell shape signature between cells in the anterior and posterior projections (Tukey’s contrasts; PP - AP *p* = 0.997 at time point 1; PP - AP *p* = 0.999 at time point 2 and PP - AP *p* = 0.896 at time point 3). These results indicate that the cell shape features included were more similar than different for cells from the structures at the third time point analysed.

Of the attributes analysed, skewness (Kruskal-Wallis test; *p* = 0.008 at time 1, *p* = 0.004 at time 2 and *p* < 0.0001 at time 3), sphericity (Kruskal-Wallis test; *p* < 0.001 at time 1, *p* = 0.00012 at time 2 and *p* < 0.001 at time 3) and surface area (Kruskal-Wallis test; *p* < 0.001 at time 1, *p* < 0.0001 at time 2 and *p* = 0.018 at time 3) described significant differences in cell shape across all the three time points analysed; cells in the endolymphatic sac were characterised by positive skewness values, smaller sphericity values and larger surface areas as compared with cells in both projections, which show negative values of skewness (Table 1 and Fig 3).

The differences in surface area are likely to be attributed to differences in sphericity between the cells in the three structures, but not in dimensions, as the transversal and longitudinal spread showed no significant differences.

## Availability and future directions

Origami will be made freely available and includes additional tools for visualising cell shape metrics from complex folding epithelia at the single-cell level. It is implemented within MATLAB (compatibility with version 2018b onwards). Instructions for installation and use are included with the software.

Our software can accept pre-segmented data, making it compatible with segmentation algorithms of the user’s choice, potentially allowing for data acquired using other 3D imaging techniques such as tomography to be analysed. Segmented data must represent cell shape accurately, and so the choice of imaging technique that can faithfully detect 3D cell shape alongside membrane or cytoplasm-based segmentation is critical.

We used *a priori* knowledge of the otic epithelium organisation to set the apico-basal axis of the epithelial sheet [2,8,39]. It is essential to know the apico-basal orientation of cells to apply Origami to any new structure studied. We also assumed that individual cells do not violate this organisation, as this cannot be detected without additional polarity-specific labels. In such a case, polarity data from our analysis can be complemented with information from polarity-specific labelling to track such behaviour. Moreover, to compute shape features oriented along an alternative axis of polarity, the pipeline can accept pre-assigned polarity as a cell-specific vector-list to compute oriented shape features.

We expect Origami to be applied to studying a wide range of morphogenetic processes and contributing to our understanding of the biomechanical processes underpinning them.

## Acknowledgements

Work was supported by a BBSRC project grant to TTW, SB and AFF (BB/M01021X/1). Imaging was carried out in the Wolfson Light Microscopy Facility at the University of Sheffield, supported by a BBSRC ALERT14 award to TTW and SB for light-sheet microscopy (BB/M012522/1). AAJ was funded by a Doctoral Training Award from the BBSRC White Rose Doctoral Training Partnership in Mechanistic Biology (BB/M011151/1). AFF is supported by the Royal Academy of Engineering Chair in Emerging Technologies Scheme (CiET1819\19) and the MedIAN Network (EP/N026993/1) funded by the Engineering and Physical Sciences Research Council (EPSRC). We thank N. van Hateren for assistance with imaging, Y. Lu for assistance with preliminary analysis, S. Burbridge and M. Marzo for technical support, and the Sheffield Aquarium Team for zebrafish husbandry.

## Supplementary Materials and Methods

### Zebrafish husbandry

All zebrafish work was reviewed and approved by the Project Applications and Amendments Committee of the University of Sheffield Animal Welfare and Ethical Review Body (AWERB). Work was performed under licence from the UK Home Office and according to recommended standard husbandry conditions [1,2]. The transgenic line used to image the cell membranes in the otic vesicle was *Tg(smad6b:mGFP)*, a gift from Robert Knight [3]. To facilitate imaging, the transgenic line was raised on a *casper* (*mitfa*^*w2/w2*^; *mpv17*^*a9/a9*^) (ZDB-GENO-080326-11) background that lacks all body pigmentation. Embryos were raised in E3 medium (5 mM NaCl, 0.17 mM KCl, 0.33 mM CaCl_2_, 0.33 mM MgSO_4_, 0.0001 % Methylene Blue). Embryonic stages are given as hours post fertilisation (hpf) at 28.5°C. For live imaging, zebrafish were anaesthetised with 0.5 mM Tricaine methylsulfonate and dechorionated.

### Microscopy

Dechorionated embryos were mounted in 0.8% Low Melting Point Agarose in E3 for microscopy. All imaging was performed at 28°C and using the 488 nm excitation laser line corresponding to the GFP membrane label. Image volume files were cropped to include the structure of interest and a small flanking region of the epithelium surrounding it.

For Airyscan confocal microscopy (Fig 1a), dechorionated embryos were mounted laterally in 0.8% Low Melting Point Agarose in E3 in the centre of a 35mm Wilco glass-bottomed Petri dish. E3 with tricaine was added to the Petri dish after the agarose had set. The image stack of 35 *z*-slices was acquired in a ZEISS LSM 880 Airyscan Confocal Microscope with a 40× objective and a *z*-step size of 1 µm.

Light-sheet microscopy (data for pipeline validation): Dechorionated embryos were mounted in agarose in a glass capillary for imaging on the Zeiss Z1 Light-sheet microscope. The microscope chamber was filled with E3 with tricaine. Image stacks varying from 60 to 110 *z*-slices depending on the age of the embryo were acquired with a 20× objective, 2.3 zoom, and a *z*-step size of 0.5 µm.

### Synthetic data generation

Synthetic images were generated in MATLAB (2018b, MathWorks) to resemble 3D volumes of folding, cell-membrane-labelled epithelia such as those in the zebrafish otic vesicle depicted in Fig 1a. Each synthetic epithelium was 160 µm x 160 µm along the epithelial plane (XY plane), consisted of about 320 individual cells and showed two projecting peaks with opposing direction of folding. The height of the peaks and the curvature of the epithelium were varied to three levels each, such that 9 individual synthetic epithelia were generated (Fig 2a).

The following function defined the surface geometry of each synthetic epithelium generated

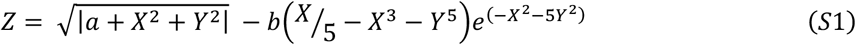

where X and Y are positions on a regular square grid (21 × 21 points) ranging from ‘-4’ to ‘4’ units with an increment of ‘0.4’ units – where each unit = 20 µm. The parameter ‘*a’* influences the radius of curvature of the epithelium (*a* = 5, 20, 80 with a resulting radius of curvature of 106 µm, 134 µm and 200 µm respectively) and ‘*b*’ controls the height of the folded peaks (*b* = 5, 10, 15 with resulting peaks of height 35 µm, 71 µm and 106 µm respectively). Centres of cells (*n* = 320) in the synthetic epithelium were initiated by randomly placing points on this surface, with a minimum distance of 8 µm between them and a padding of 8 µm from the edge of the grid. The resulting set of points were nearly equally spaced.

To convert these surfaces into image volumes, the cell centre positions were then resampled to a volume of isotropic resolution with pixel size of 0.2 µm, resulting in 800 pixels x 800 pixels x >800 pixels, since the *z* dimension was adjusted to accommodate cell positions spanning more than 800 pixels. A Voronoi diagram was generated from the resampled cell centres. The edges of the Voronoi cells were extended 5 µm (26 pixels) orthogonal to the epithelial surface to set cell height and 0.4 µm (2 pixels) in the epithelial plane to set cell membrane thickness. These extended Voronoi edges were used to define a 3D network of polygons as cell membranes. Pixels on the grid that lay within the cell membrane polygons were assigned an intensity value of ‘1’.

The synthetic images generated were then convolved with a Gaussian PSF using a Fast Fourier Transform (FFT)-based convolution (FFT-based convolution; Bruno Luong, MathWorks File Exchange, accessed Oct 2020) to resemble real-world imaging conditions. The PSF was simulated using the PSF Generator plugin in Fiji [4,5], assuming the following experimental parameters: Numerical Aperture of collection objective lens = 0.5, wavelength of illumination = 532 nm, voxel size = 0.2 µm x 0.2 µm x 0.2 µm. The resulting full width at half maximum (FWHM) of the PSF was 0.6 µm x 0.6 µm x 0.8 µm (3 pixels x 3 pixels x 4 pixels). Finally, after combining each of the images (*n* = 9) with the three levels of Gaussian and Poisson noise using the ‘imnoise’ function in MATLAB, 27 synthetic image volumes were generated for performing the validation tests. Ground truth to assess segmentation quality was produced from the 9 uncorrupted image volumes.

Polarity ground truth for the synthetic dataset was generated by producing surface normals to the surface functions described by equation (S1), using the SurfNorm function in MATLAB (version 2018b; MathWorks, Natick MA, US).

### Membrane-based segmentation

The parameters used to segment our datasets in ACME were different for the synthetic dataset and the real light-sheet data in part, due to differences in voxel resolution (0.2 µm x 0.2 µm x 0.2 µm for the synthetic dataset and 0.1 µm x 0.1 µm x 0.5 µm for the light-sheet data). These parameters were as follows;

#### For synthetic epithelia

1. Radius of median filter for denoising – 3.0 pixels (image volumes with noise level 2 and 3), 2.0 pixels (noise level 1)

2. Resampling ratio – 2.5, 2.5, 2.5 (all image volumes)

3. neighbourhood size for membrane signal enhancement filter – 2.0 (noise level 1 and 2), 3.0 (noise level 3)

4. neighbourhood size for Tensor voting – 1.0 (all image volumes)

5. watershed segmentation threshold – 2.0 (noise level 1 and 2), 3.0 (noise level 3)

#### For fluorescence *in-vivo* data

1. Radius of median filter for denoising – 0.3 pixels

2. Resampling ratio – 2, 2, 0.39 (resampling to isotropic voxel resolution)

3. neighbourhood size for membrane signal enhancement filter – 0.7

4. neighbourhood size for Tensor voting – 1.0

5. watershed segmentation threshold – 2.0

### Classifying cells

Cells were classified as lying within the folding structure or the neighbouring epithelium by clustering the centroids of the segmented cells by the mean curvature (Fig S1), that is, the average of the principal curvatures at each vertex [6,7] of the surface mesh generated in the first part of the Origami pipeline. The mean curvature values showed a bimodal distribution, which could be resolved into a population of points on the folding structure and another consisting of points on the neighbouring non-folding epithelium. Cells at the edges of the image volume were discarded to avoid broken cells.

**Fig S1:**
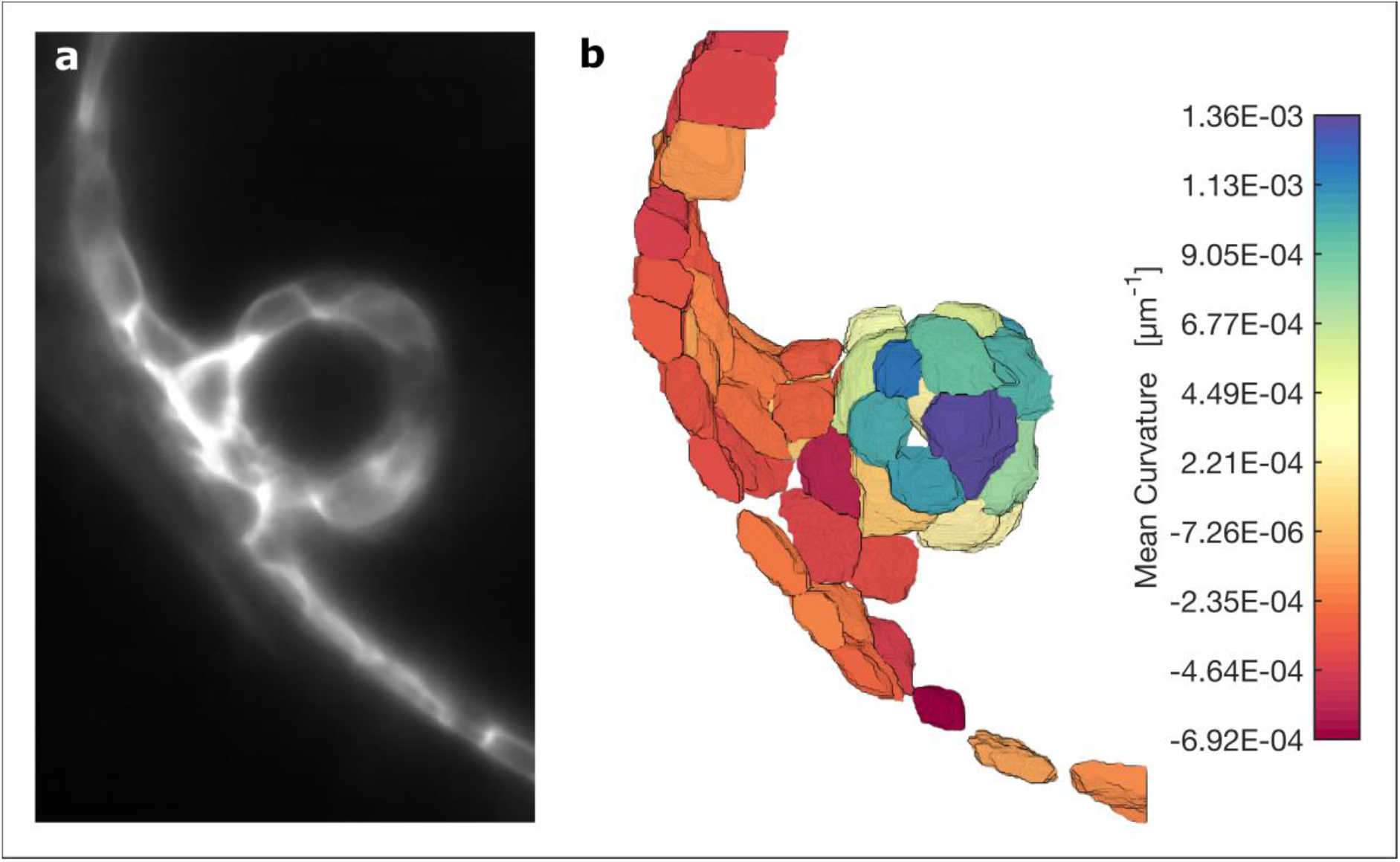
Cell-specific mean curvature of epithelium. **a**. Single slice through a light-sheet image volume of a region around an anterior projection in the otic vesicle of a 50.5 hpf zebrafish embryo. **b**. 3D rendering of segmented cells from the same region with individual cells assigned colour values corresponding to the mean curvature at the apical surface of the cell.

